# NeurotoxKb: compilation, curation and exploration of a knowledgebase of environmental neurotoxicants specific to mammals

**DOI:** 10.1101/2021.01.12.426435

**Authors:** Janani Ravichandran, Bagavathy Shanmugam Karthikeyan, Palak Singla, S. R. Aparna, Areejit Samal

**Author notes:** J.R. and B.S.K. contributed equally to this work and should be considered as Joint-First authors. Corresponding author (A. Samal), **Address for correspondence:** Areejit Samal, Computational Biology Group, The Institute of Mathematical Sciences (IMSc), CIT Campus, Taramani, Chennai 600113 India.

## Abstract

Exposure to environmental neurotoxicants is a significant concern due to their potential to cause permanent or irreversible damage to the human nervous system. Here, we present the first dedicated knowledgebase, NeurotoxKb, on environmental neurotoxicants specific to mammals. Using a detailed workflow, we have compiled 475 potential non-biogenic neurotoxicants from 835 published studies with evidence of neurotoxicity specific to mammals. A unique feature of NeurotoxKb is the manual curation effort to compile and standardize the observed neurotoxic effects for the potential neurotoxicants from 835 published studies. For the 475 potential neurotoxicants, we have compiled diverse information such as chemical structures, environmental sources, chemical classification, physicochemical properties, molecular descriptors, predicted ADMET properties, and target human genes. To better understand the prospect of human exposure, we have explored the presence of potential neurotoxicants in external exposomes via two different analyses. By analyzing 55 chemical lists representing global regulations and guidelines, we reveal potential neurotoxicants both in regular use and produced in high volume. By analyzing human biospecimens, we reveal potential neurotoxicants detected in them. Lastly, a construction of the chemical similarity network and ensuing analysis revealed the diversity of the toxicological space of 475 potential neurotoxicants. NeurotoxKb is accessible online at: https://cb.imsc.res.in/neurotoxkb/.

## 1. Introduction

The human nervous system is both complex and sensitive to environmental exposures [1,2] Exposure to environmental chemicals can lead to various neurological disorders and neurotoxic effects which can manifest at any stage of human life, from infancy to old age [3,4]. The common sources of daily exposure to these environmental chemicals include industrial chemicals, agrochemicals, exhausts from automobile or industries, personal care products, and household supplies [5,6]. Exposures to these environmental neurotoxicants are of significant concern as they can cause permanent or irreversible damage to the nervous system [7,2].

In the last few decades, several studies have documented the neurotoxic effects of heavy metals such as arsenic, lead, manganese and mercury, and other groups of environmental chemicals such as Polychlorinated biphenyls (PCBs), Perfluoroalkylated substances (PFAS) and Organotins [8,9,3,4,6]. In comparison to chemicals tested for neurotoxicity so far, the space of chemicals in commerce is huge. Specifically, there are over 100000 chemicals in commerce in the European Union (EU) and USA, and only a tiny fraction of them have been tested for neurotoxicity to date [8,10]. Some reasons for this gap in current knowledge on environmental neurotoxicants include the lack of systematic testing methods for neurotoxicity and the inherent complexity of neurotoxicological assessments [11,8,6]. Despite these limitations, there have been some efforts to compile potential neurotoxicants with evidence specific to mammals from published literature [12,8,9,13,14].

To the best of our knowledge, there have been four previous efforts to compile environmental neurotoxicants with evidence specific to mammals from published literature [12,8,9,13,14]. In 1976, the US Environmental Protection Agency (EPA) published an extensive report on chemicals tested for neurotoxicity [12]. Since 2006, Grandjean and Landrigan have compiled 214 developmental neurotoxicants based on published human-specific studies documented in toxicological resources [8,9]. In 2015, Mundy *et al* [13] compiled a list of 97 potential neurotoxicants demonstrating effects on neurodevelopment, and thereafter, in 2017, Aschner *et al* [14] compiled a list of 33 potential neurotoxicants triggering developmental neurotoxicity *in vivo*. Although the lists of potential neurotoxicants compiled by Grandjean and Landrigan [8], Mundy *et al* [13], and Aschner *et al* [14] are available via the CompTox dashboard [15], there is no dedicated online resource to date on environmental neurotoxicants. Here, we address this unmet need by building the first dedicated online knowledgebase on potential non-biogenic neurotoxicants with published evidence specific to mammals.

To create the environmental <u>Neurotox</u>icants <u>K</u>nowledge<u>b</u>ase NeurotoxKb, we have developed a detailed workflow (Figure 1) to identify potential non-biogenic neurotoxicants with evidence specific to mammals from published literature. An important limitation of the existing resources on neurotoxicants is in their compilation of observed neurotoxic effects using non-standardized vocabulary [12–14] or the complete lack thereof [8,9]. To overcome this limitation of existing resources, we have performed an extensive manual curation effort to compile, unify and standardize the reported neurotoxic effects for potential neurotoxicants in published literature, into standardized neurotoxic endpoints. In a nutshell, we have identified here 475 potential neurotoxicants which are non-biogenic and have evidence of neurotoxicity specific to mammals from published studies. For these 475 potential neurotoxicants, our compilation includes observed neurotoxic effects in terms of 148 standardized neurotoxic endpoints curated from 835 published studies specific to mammals. For the 475 potential neurotoxicants, we have compiled additional information including chemical structures, chemical classification, environmental sources, physicochemical properties, predicted ADMET properties, molecular descriptors and target human genes. The entire information compiled in NeurotoxKb, on the 475 potential neurotoxicants specific to mammals, is accessible at: https://cb.imsc.res.in/neurotoxkb.

**Figure 1:**
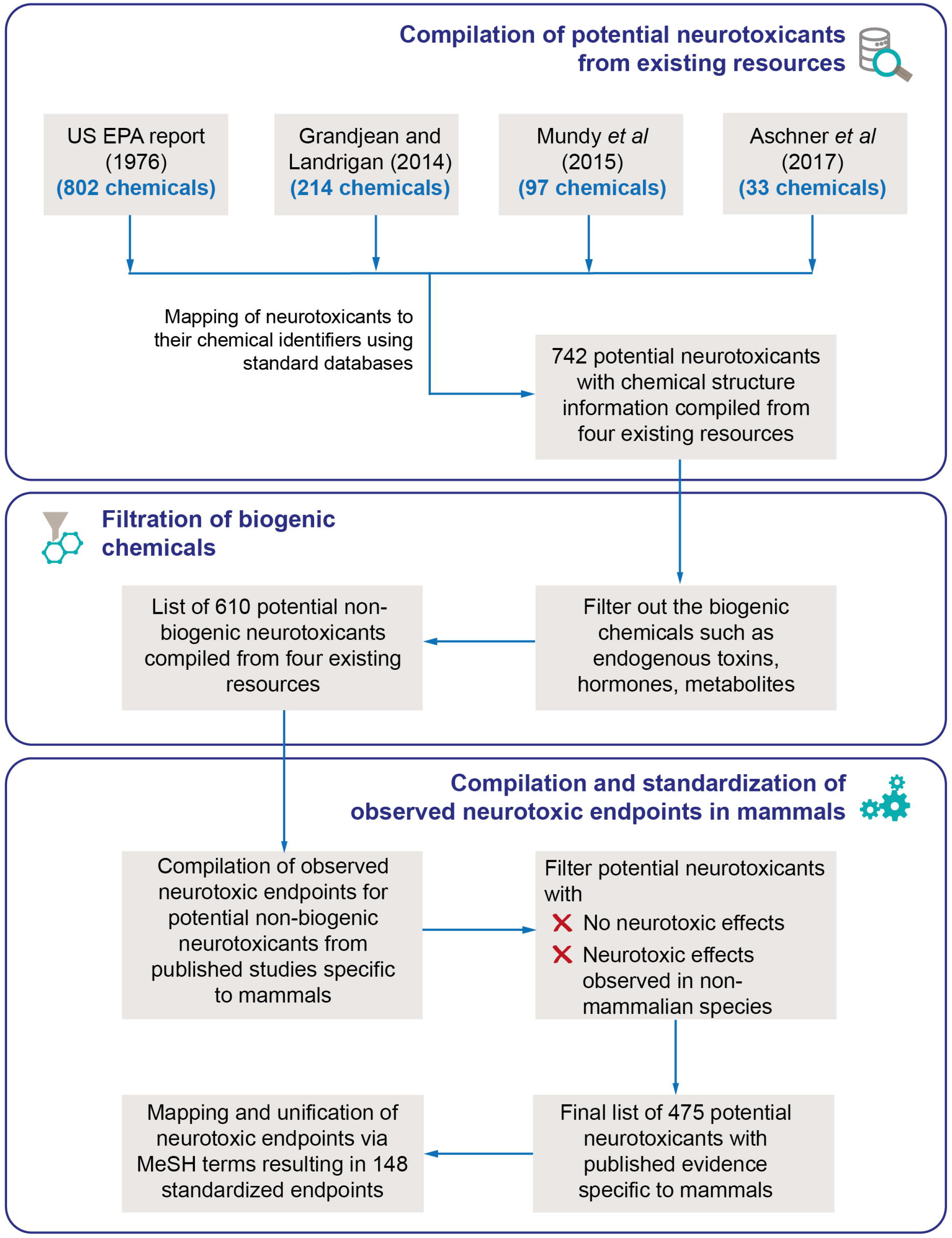
Schematic workflow describing the compilation of 475 potential non-biogenic neurotoxicants along with published evidence of observed neurotoxic endpoints specific to mammals.

Human exposure to neurotoxicants can occur via diverse environmental sources [5,6]. Hence, it is important to understand the current state of regulation and monitoring of environmental neurotoxicants through the perspective of exposomes. To this end, we have analyzed the presence of potential neurotoxicants across 55 chemical lists which include inventories, regulations and guidelines. Notably, based on the source or route of exposure, we have classified these 55 chemical lists into different categories of exposome. Thus, the presence of neurotoxicants in these 55 chemical lists is a clear indication of their presence in human exposome. As detection of environmental chemicals in biospecimens is a proof of their exposure, we have also analyzed the presence of potential neurotoxicants among chemicals detected in different human biospecimens such as blood, urine, placenta and human milk [16–19]. Furthermore, based on comparative analyses with current chemical regulations and guidelines, we present a hazard priority list of 18 potential neurotoxicants. In short, we show the utility of our resource in aiding regulatory bodies worldwide in prioritization of hazardous chemicals, to streamline their monitoring and regulation.

Since neurotoxicants affect the nervous system via diverse molecular mechanisms [6], a network biology [20] perspective on associations between potential neurotoxicants and their target human genes or proteins can shed insights on molecular mechanisms behind observed neurotoxic effects. Therefore, we have constructed and analyzed a bipartite network of potential neurotoxicants in NeurotoxKb and their target human neuroreceptors. Moreover, there is growing interest in chemical structure based prioritization approaches in risk assessment and toxicology [21,22]. Hence, we have also performed a similarity based analysis of the chemical space of potential neurotoxicants. By constructing a chemical similarity network, we show that the space of potential neurotoxicants in NeurotoxKb is highly diverse. Overall, NeurotoxKb is a comprehensive knowledgebase on potential environmental neurotoxicants specific to mammals which will enable future research in neurotoxicology.

## 2. Methods

### 2.1. Building a knowledgebase of environmental neurotoxicants specific to mammals

#### 2.1.1. Compilation and filtration of potential non-biogenic neurotoxicants from existing resources

We started building the curated knowledgebase on environmental neurotoxicants, namely NeurotoxKb, with experimental evidence on neurotoxicity specific to mammals, by compiling potential neurotoxicants from four existing resources [12,8,9,13,14] in published literature (Figure 1; Table 1).

**Table 1:** Comparison of the features including compiled information captured in NeurotoxKb for the potential neurotoxicants with respect to four existing resources.

| Feature | NeurotoxKb | US EPA report (1976) | Grandjean and Landrigan (2014) | Mundy *et al* (2015) | Aschner *et al* (2017) |
| --- | --- | --- | --- | --- | --- |
| Number of potential neurotoxicants | 475 | 802 | 214 | 97 | 33 |
| Web interface | Yes | No | No | Yes via CompTox | Yes via CompTox |
| Compilation of neurotoxic endpoints | Yes | Yes | No | Yes | Yes |
| Standardization of neurotoxic endpoints | Yes | No | No | No | No |
| Classification based on environmental source | Yes | No | Yes | Yes | Yes |
| Classification based on chemical structure | Yes | No | No | No | No |
| Presence in chemical regulation or guideline | Yes | No | No | Yes | Yes |
| Information on external exposomes | Yes | No | No | No | No |
| Presence in human biospecimen | Yes | No | No | No | No |
| Chemical identifiers | PubChem or CAS or MeSH | CAS | CAS | DSSTox substance identifier or CAS | DSSTox substance identifier or CAS |
| Download of 2D structure | SDF, MOL, MOL2 | No | No | MOL | MOL |
| Download of 3D structure | SDF, MOL, MOL2, PDB, PDBQT | No | No | No | No |
| Physicochemical properties | Yes | No | No | Yes | Yes |
| Molecular descriptors | Yes | No | No | No | No |
| Predicted ADMET properties | Yes | No | No | Yes | Yes |
| Chemical-gene association | Yes | No | No | Yes | Yes |
| Chemical similarity filter | Yes | No | No | No | No |

Firstly, we considered 802 potential neurotoxicants compiled in the US Environmental Protection Agency (EPA) report [12] published in 1976 on neurotoxic chemicals. From published literature, the US EPA report had compiled 802 chemicals tested for neurotoxic effects upon exposure on various living organisms including mammals and non-mammals [12]. Secondly, we have considered 214 potential neurotoxicants compiled by Grandjean and Landrigan [8,9] to which humans are vulnerable upon exposure in early stages of development. For compiling their list, Grandjean and Landrigan [8,9] had employed PubMed literature mining and toxicological resources such as TOXNET [23], TOXLINE [24] and Hazardous Substances Data Bank (HSDB) [25]. Note that Grandjean and Landrigan had first published a list of 201 potential human neurotoxicants in 2006 [8] which was subsequently expanded to 214 potential human neurotoxicants by them in 2014 [9]. Thirdly, we have considered the 97 potential neurotoxicants compiled by Mundy *et al* [13] (https://comptox.epa.gov/dashboard/chemical_lists/DNTEFFECTS) that have demonstrated effects on neurodevelopment. Fourthly, we have considered the 33 potential neurotoxicants compiled by Aschner *et al* [14] (https://comptox.epa.gov/dashboard/chemical_lists/DNTINVIVO) that have evidence of triggering developmental neurotoxicity *in vivo*.

We remark that three of the above-mentioned four lists of potential neurotoxicants considered here (Figure 1), namely Grandjean and Landrigan [8], Mundy *et al* [13] and Aschner *et al* [14], are among the six lists of potential neurotoxicants captured by the CompTox chemistry dashboard [15] (https://comptox.epa.gov/dashboard/chemical_lists). Since our aim is to compile potential neurotoxicants specific to mammals, we have not considered three other lists of potential neurotoxicants captured by the CompTox chemistry dashboard. Specifically, we have not considered the list ‘DNT Screening Library’ (https://comptox.epa.gov/dashboard/chemical_lists/DNTSCREEN) that compiles potential neurotoxicants with experimental evidence specific to Zebrafish. Similarly, we have not considered the two lists, namely ‘Neurotoxicants from PubMed’ (https://comptox.epa.gov/dashboard/chemical_lists/LITMINEDNEURO) and ‘Neurotoxicants Collection from Public Resources’ (https://comptox.epa.gov/dashboard/chemical_lists/NEUROTOXINS), as both lists gather potential neurotoxicants from literature without compiling information on the test organisms for neurotoxicity.

Next, we mapped the 802, 214, 97 and 33 potential neurotoxicants compiled from the US EPA report [12], Grandjean and Landrigan [9], Mundy *et al* [13] and Aschner *et al* [14], respectively, to chemical identifiers in standard databases such as PubChem [26], Chemical Abstract Service (CAS; https://www.cas.org/) and the Comparative Toxicogenomics Database (CTD) [18]. While mapping the potential neurotoxicants to their chemical structure, we have removed any potential neurotoxicant in the four lists that could not be mapped to a chemical identifier or represents a chemical mixture rather than individual chemical entity. This resulted in a non-redundant list of 742 potential neurotoxicants compiled from the four above-mentioned resources (Figure 1).

Next, we have removed any chemical from the non-redundant list of 742 potential neurotoxicants compiled from the four above-mentioned resources that are of biological origin such as snake venoms, plant or microbial toxins, and hormones. This removal of potential biogenic neurotoxins is motivated by our exclusive focus on man-made environmental neurotoxicants. This resulted in a list of 610 potential non-biogenic neurotoxicants compiled from the four above-mentioned resources (Figure 1).

In summary, we have compiled from four existing resources, a curated list of 610 potential non-biogenic neurotoxicants along with their two-dimensional (2D) and three-dimensional (3D) chemical structure information via the above-mentioned steps in our workflow (Figure 1).

#### 2.1.2. Compilation and standardization of observed neurotoxic endpoints for environmental neurotoxicants specific to mammals

In order to develop a comprehensive resource on environmental neurotoxicants, it is necessary to compile the observed neurotoxic endpoints (or adverse effects) upon exposure to neurotoxicants from the published literature. Although the four existing resources [9,12–14] on potential neurotoxicants considered here compile observed neurotoxic endpoints upon chemical exposure, a lack of standardization in reporting of the adverse effects across the resources limit their utility for toxicological risk assessment. To address this unmet need and enable future research in neurotoxicity, we next compiled and manually curated the observed neurotoxic endpoints for the 610 potential non-biogenic neurotoxicants identified via the above-mentioned steps in our workflow (Figure 1).

Firstly, we have compiled from the USA EPA report [12], the observed neurotoxic endpoints for potential non-biogenic neurotoxicants along with the information on test organisms including mammals and non-mammals in the published experimental studies. Note that the USA EPA report [12] also compiles observations of no neurotoxic effects for potential neurotoxicants from published experimental studies.

Secondly, Mundy *et al* [13] and Aschner *et al* [14] have compiled potential developmental neurotoxicants along with the information on their observed neurotoxic endpoints from published experimental studies in rodents and primates. However, the compilation of neurotoxic endpoints in Mundy *et al* [13] and Aschner *et al* [14] is much less detailed in comparison to the USA EPA report [12]. Specifically, Mundy *et al* [13] have reported the neurotoxic endpoints from published studies after their broad categorization into 3 terms, namely, behaviour, morphology, and neurochemistry. Similarly, Aschner *et al* [14] have reported the neurotoxic endpoints from published studies after their broad categorization into 40 terms. However, we believe that a detailed compilation of neurotoxic endpoints for potential neurotoxicants from published studies specific to mammals can render a valuable toxicological resource that can aid in early identification and regulation of hazardous chemicals. Therefore, we have performed a manual curation of the 287 published studies compiled by Mundy *et al* [13] and Aschner *et al* [14] to collect detailed neurotoxic endpoints for potential non-biogenic neurotoxicants covered by the two resources.

Thirdly, Grandjean and Landrigan [8,9] have compiled a list of chemicals potentially toxic to the human nervous system from published literature. However, Grandjean and Landrigan [8,9] have not compiled the observed neurotoxic endpoints for the potential neurotoxicants from associated published literature. Therefore, we have performed an extensive manual curation effort to compile the observed neurotoxic effects specific to humans from HSDB [25] for the potential neurotoxicants in the list by Grandjean and Landrigan [8,9]. Note that HSDB [25] (which has been integrated into PubChem [26]) was used by Grandjean and Landrigan [8,9] to compile their list of 214 potential human neurotoxicants. During this manual curation effort, we were unable to gather experimental evidence specific to mammals from HSDB [25] for some of the 214 potential human neurotoxicants in the list by Grandjean and Landrigan [8,9]. For such potential neurotoxicants in the list by Grandjean and Landrigan [8,9] without any documented evidence of neurotoxicity in HSDB [25], we have performed additional literature search to gather any published evidence of neurotoxicity specific to mammals.

At the end of the above-mentioned steps to compile observed neurotoxic endpoints specific to mammals for 610 potential non-biogenic neurotoxicants from existing resources [9,12–14], HSDB [25] and published literature, we were able to gather published experimental evidence specific to mammals for only 475 out of 610 potential non-biogenic neurotoxicants (Figure 1; Supplementary Table S1). These 475 potential non-biogenic neurotoxicants with experimental evidence specific to mammals from 835 published articles have been compiled in our environmental <u>Neurotox</u>icants <u>K</u>nowledge<u>b</u>ase, namely NeurotoxKb, which is accessible at: http://cb.imsc.res.in/neurotoxkb.

Finally, we undertook an extensive manual curation effort to standardize the compiled information on detailed neurotoxic effects observed in 835 published studies specific to mammals for the 475 potential non-biogenic neurotoxicants in NeurotoxKb. For the unification and standardization of this compiled information on neurotoxic effects of the 475 potential neurotoxicants, we have leveraged Medical Subject Headings (MeSH) terms [27]. Through this exercise, we were able to map, unify and standardize a compiled list of 900 terms referring to observed neurotoxic effects from 835 published studies on 475 potential neurotoxicants to 148 standardized neurotoxic endpoints based on MeSH terms (Figure 1; Supplementary Table S2).

### 2.2. Classification of neurotoxicants based on environmental source or chemical structure

Information on the major sources of exposure is vital for chemical regulation and monitoring by agencies. Therefore, we have compiled the environmental sources for the 475 potential neurotoxicants in NeurotoxKb. Specifically, NeurotoxKb has classified the 475 potential neurotoxicants into 6 broad categories of environmental sources, namely, Agriculture and Farming, Consumer Products, Industry, Intermediates, Medicine and Healthcare, and Pollutant, and 41 sub-categories (Figure 2).

**Figure 2:**
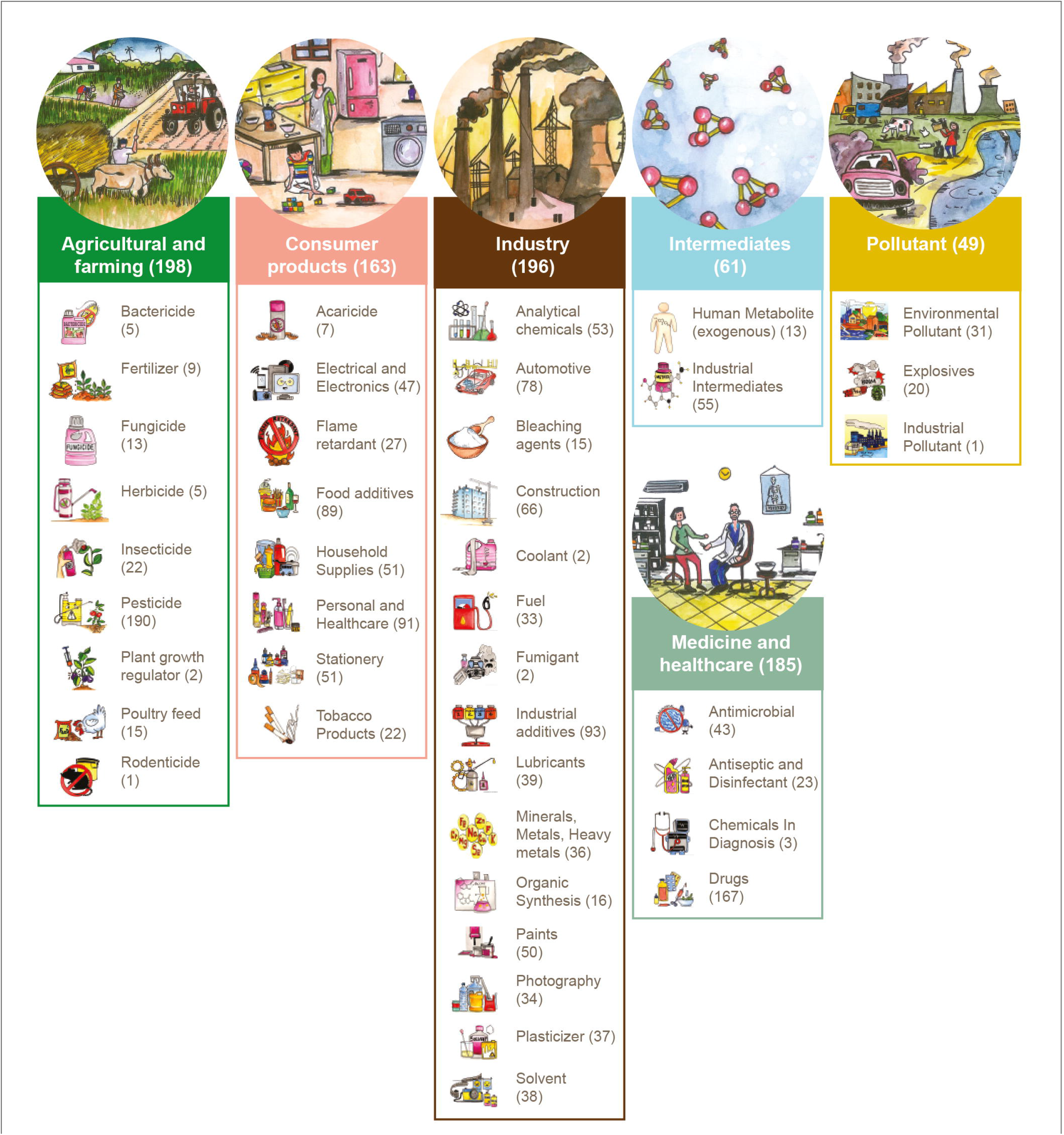
Classification of the 475 potential neurotoxicants in NeurotoxKb into 6 broad categories and 41 sub-categories based on their environmental source. The number of potential neurotoxicants in each category or sub-category is mentioned besides the category or sub-category within parenthesis. Note that a potential neurotoxicant can belong to more than one category or sub-category of environmental sources.

Furthermore, we have also classified the 475 potential neurotoxicants in NeurotoxKb based on their chemical structure. Specifically, we have employed ClassyFire [28] (http://classyfire.wishartlab.com/) for a hierarchical chemical classification into kingdom, superclass, class and subclass, of the 475 potential neurotoxicants in NeurotoxKb. Note that information on chemical class of potential neurotoxicants can be used to draw inferences on their nature and behaviour.

### 2.3. Physicochemical and ADMET properties of neurotoxicants

We have used cheminformatics software to compile physicochemical properties, molecular descriptors and predicted Absorption, Distribution, Metabolism, Excretion and Toxicity (ADMET) properties for the 475 potential neurotoxicants in NeurotoxKb. This information will assist both computational and experimental research on neurotoxicants in future. The physicochemical properties and the molecular descriptors for the 475 potential neurotoxicants were computed using RDKit (https://www.rdkit.org/), PaDEL [29] and Pybel [30]. The ADMET properties for the 475 potential neurotoxicants were predicted using admetSAR 2.0 [31], pkCSM [32], SwissADME [33], Toxtree 2.6.1 [34] and vNN server [35].

### 2.4. Target genes of environmental neurotoxicants

Identification of target human genes or proteins of environmental neurotoxicants can shed light on complex molecular mechanisms via which these chemicals cause neurotoxicity. We have used ToxCast [36] to identify the target human genes or proteins of the 475 potential neurotoxicants in NeurotoxKb. To retrieve the list of target human genes perturbed by potential neurotoxicants, we have used ToxCast invitroDB3 dataset released in August 2019 [37]. The assay summary information file (Assay_Summary_190708.csv) of the ToxCast invitroDB3 dataset [37] contains detailed annotation for the different ToxCast assays. Using this assay summary information file, we have identified human-specific ToxCast assays and their corresponding endpoints. If a tested chemical in a human-specific ToxCast assay is found to be ‘active’ for the corresponding assay component endpoint, then the corresponding gene or protein is assigned as a target of the chemical.

### 2.5. Web interface of NeurotoxKb

The web interface of NeurotoxKb (Figure 3) has been created using Bootstrap 4, CSS, HTML, jQuery (https://jquery.com/) and PHP (http://php.net/). The compiled database on the 475 potential neurotoxicants is stored and retrieved using MariaDB (https://mariadb.org/) and Structured Query Language (SQL), respectively. Interactive visualization of the compiled information in NeurotoxKb is facilitated by Cytoscape.js (http://js.cytoscape.org/), Google Charts (https://developers.google.com/chart/) and Plotly (https://plotly.com/javascript/). NeurotoxKb is hosted on an Apache (https://httpd.apache.org/) web server running on Debian 9.4 Linux Operating System.

**Figure 3:**
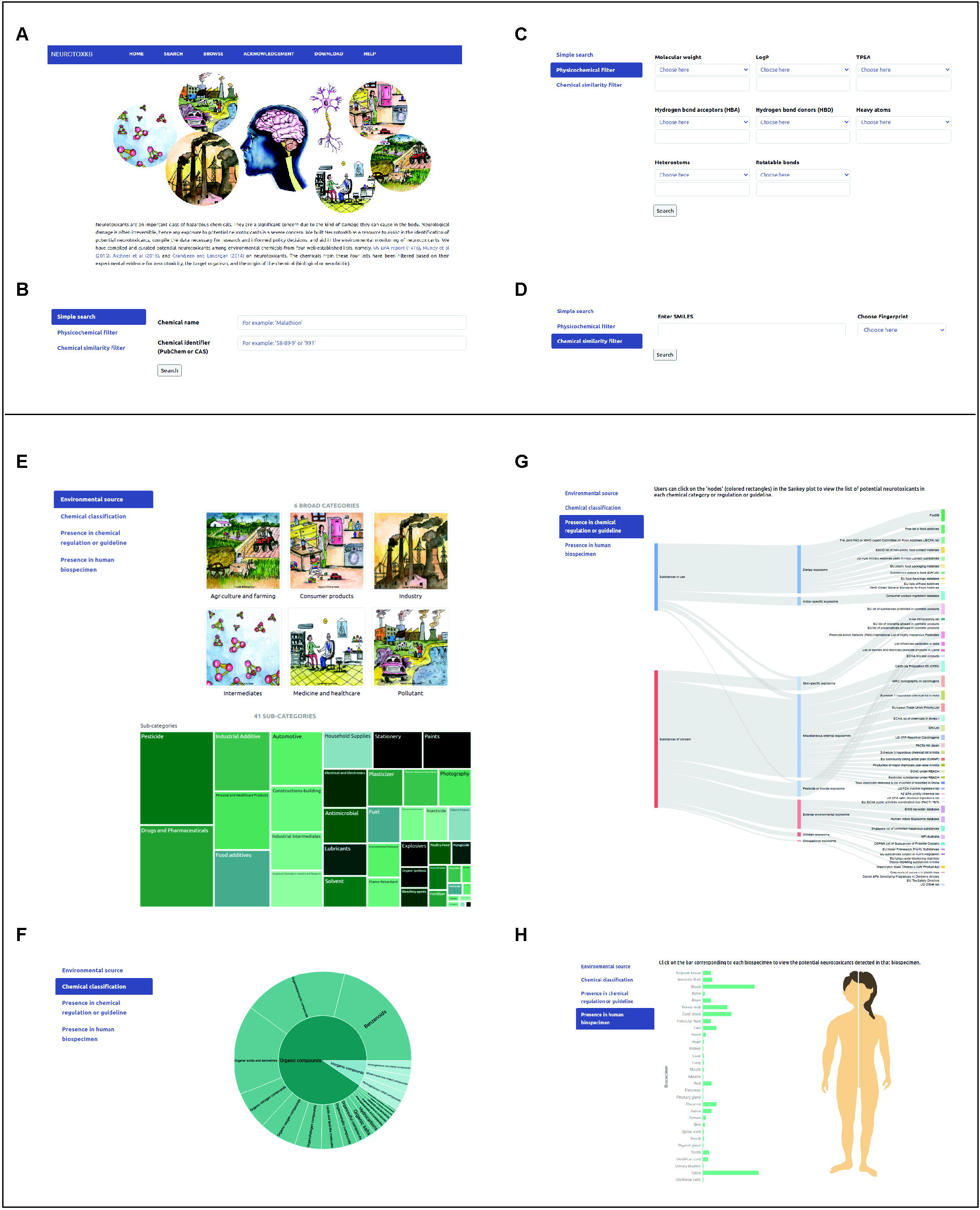
The web interface of NeurotoxKb. **(A)** The screenshot displays the home page of NeurotoxKb. NeurotoxKb has options to search and retrieve information on potential neurotoxicants, namely, **(B)** Simple search to retrieve potential neurotoxicants using their chemical names or identifiers, **(C)** Physicochemical filter to retrieve potential neurotoxicants based on their physicochemical properties, and **(D)** Chemical similarity filter to retrieve potential neurotoxicants that are structurally similar to a query compound. NeurotoxKb also has options to browse information on potential neurotoxicants based on their **(E)** Environmental source classification, **(F)** Chemical classification, **(G)** Presence in chemical regulation or guideline, and **(H)** Presence in human biospecimen.

### 2.6. Exploration of potential neurotoxicants across chemical inventories, regulations and guidelines

Understanding the environmental sources and routes of exposure to neurotoxicants will be critical for monitoring and mitigation of their adverse effects on humankind. We have explored the presence of neurotoxicants in external exposomes via a comparative analysis with 55 publicly available chemical lists including inventories, regulations and guidelines (Figure 4; Supplementary Table S3). These 55 chemical lists were broadly classified into two categories, namely ‘Substances in use (SIU)’ and ‘Substances of concern (SOC)’ (Figure 4; Supplementary Table S3). SIU lists consist of chemicals that are permitted or found to be in regular use while SOC lists consist of chemicals that are marked hazardous, regulated or restricted by government or independent bodies across the world [38]. Based on the source or route of human exposure, the 55 chemical lists have further been classified into 8 categories of exposomes, namely, ‘Childrens’ exposome’, ‘Dietary exposome’, ‘External environmental exposome’, ‘Indoor-specific exposome’, ‘Occupational exposome’, ‘Pesticide/biocide exposome’, ‘Skin-specific exposome’ and ‘Miscellaneous external exposome’ (Figure 4; Supplementary Table S3), and these contribute to the total external exposome of humans. Here, we have performed a comparative analysis for potential neurotoxicants with SIU and SOC lists similar to that performed for potential endocrine disruptors in our recent publication [38]. Note that the presence of any potential neurotoxicant in SIU or SOC lists reflects its potential for human exposure.

**Figure 4:**
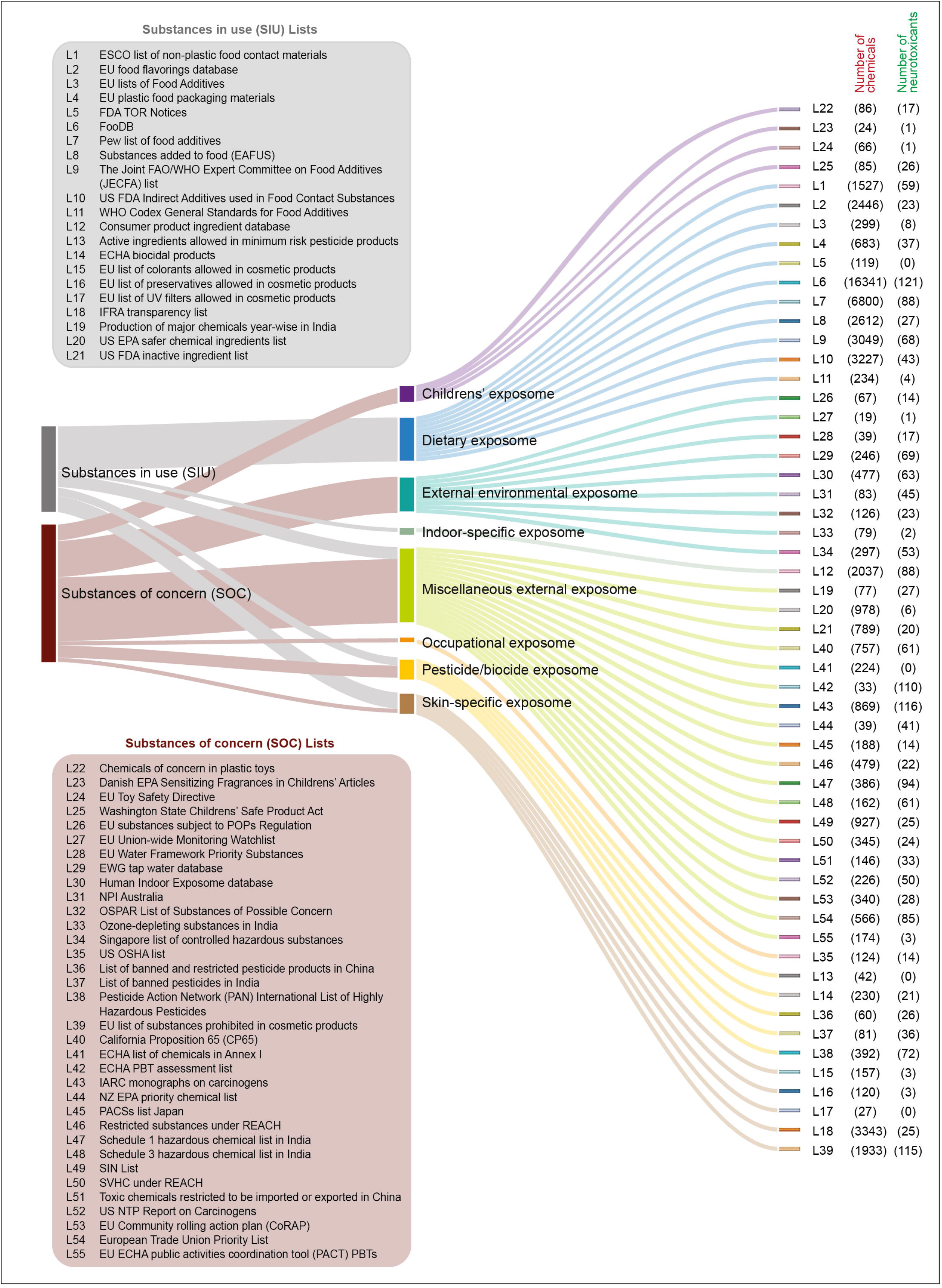
Sankey plot displays the 55 chemical lists considered for comparative analysis that are a part of chemical inventories, regulations and guidelines. These lists were broadly classified into two categories, namely, Substances in use (SIU) and Substances of concern (SOC), based on the nature of substances. Further, these lists have also been classified into 8 categories of exposome, namely, Childrens’ exposome, Dietary exposome, External environmental exposome, Indoor-specific exposome, Miscellaneous external exposome, Occupational exposome, Pesticide/biocide exposome, and Skin-specific exposome, based on the route or source of exposure. Besides each chemical list, the total number of chemicals and the number of potential neurotoxicants present in that list are shown within parenthesis.

Previously, Grandjean and Landrigan [8,9] have highlighted that several of the commercial chemicals which are produced in high volume across the world, have not been tested for their neurotoxic potential. To better understand the scale of possible human exposure to neurotoxicants, we have also explored the presence of 475 potential neurotoxicants in two publicly available lists of chemicals produced in high volume, namely, the United States High Production Volume (USHPV) database and the Organisation for Economic Cooperation and Development High Production Volume (OECD HPV) list which was last updated in 2004.

### 2.7. Exploration of potential neurotoxicants in human biospecimens

Exposome refers to the totality of exposure during the lifetime of an individual and their associated health effects [39–42]. Note that the presence of any potential neurotoxicant in a human biospecimen presents conclusive proof of human exposure and is also indicative of its potential to affect the nervous system. Here, we have explored the presence of 475 potential neurotoxicants in human biospecimens using compiled data in two resources, namely, the Exposome-Explorer [16] and CTD [18]. Using literature mining, Exposome-Explorer [16] has compiled information on environmental chemicals detected in different human biospecimens from published literature based on dietary and pollution exposures. Similarly, ‘Exposure – study associations’ in CTD [18] can be used to retrieve compiled information from published literature on environmental chemicals detected in different human biospecimens. Importantly, we have compiled information on environmental chemicals detected in different human biospecimens from two resources [16,18], and the annotation of the human biospecimens is not uniform across Exposome-Explorer [16] and CTD [18]. Therefore, we have manually curated and unified the different human biospecimens captured in the two resources, Exposome-Explorer [16] and CTD [18], into 31 different types or exposomes (Figure 2; Supplementary Table S6). For example, we have grouped the human biospecimens such as plasma, serum, blood proteins or blood cells into a single type ‘blood’ exposome in this work.

### 2.8. Chemical similarity network of environmental neurotoxicants

Chemical similarity networks (CSNs) can shed insights on the extent of scaffold diversity in the associated chemical space [43,21,44,22]. Further, chemical similarity approaches can aid in early identification of toxic chemicals [21,22] including potential neurotoxicants. To construct the CSN of neurotoxicants, we have employed the similarity metric Tanimoto coefficient (Tc) [45]. For any pair of chemicals, Tc has a value in the range 0 to 1, wherein the level of chemical similarity between two molecules is directly proportional to the corresponding Tc value. The computation of Tc between pairs of chemicals can depend on the choice of chemical fingerprints used to represent the molecules. Here, we have chosen Extended Circular Fingerprints (ECFP4) [46] while computing Tc between different pairs of potential neurotoxicants. In the CSN of potential neurotoxicants in NeurotoxKb, there are 475 nodes corresponding to the 475 potential neurotoxicants, and there is an edge between any pair of nodes if the corresponding Tc value is ≥ 0.5. The chosen cut off of Tc ≥ 0.5 to decide on significant structural similarity between pairs of chemicals was motivated by similar choice made in previous studies [47–49].

## 3. Results and Discussion

### 3.1. NeurotoxKb: Knowledgebase of environmental neurotoxicants specific to mammals

Following the workflow shown in Figure 1, we have built a manually curated knowledgebase of environmental neurotoxicants, namely NeurotoxKb, which compiles information on 475 potential non-biogenic neurotoxicants with published evidence of observed neurotoxic effects specific to mammals (Methods). NeurotoxKb is accessible at: https://cb.imsc.res.in/neurotoxkb.

Importantly, we have curated the information on potential neurotoxicants from published studies compiled in four existing resources, namely, the US EPA report [12], Grandjean and Landrigan [9], Mundy *et al* [13] and Aschner *et al* [14]. Supplementary Table S1 gives the curated list of 475 potential non-biogenic neurotoxicants in NeurotoxKb. Of these 475 potential neurotoxicants, the US EPA report [12], Grandjean and Landrigan [9], Mundy *et al* [13] and Aschner *et al* [14] capture 292, 178, 88 and 26 potential neurotoxicants, respectively, with published evidence specific to mammals (Figure 5A). Notably, among the four existing resources, the US EPA report [12] contributes a unique set of 231 out of the 475 potential neurotoxicants (~50%) compiled in NeurotoxKb with published evidence specific to mammals (Figure 5A). In other words, almost 50% of the potential neurotoxicants specific to mammals in NeurotoxKb were solely identified due to our extensive manual effort to digitize, compile, curate and organize the vast information on potential neurotoxicants captured in the US EPA report [12] published in 1976.

**Figure 5:**
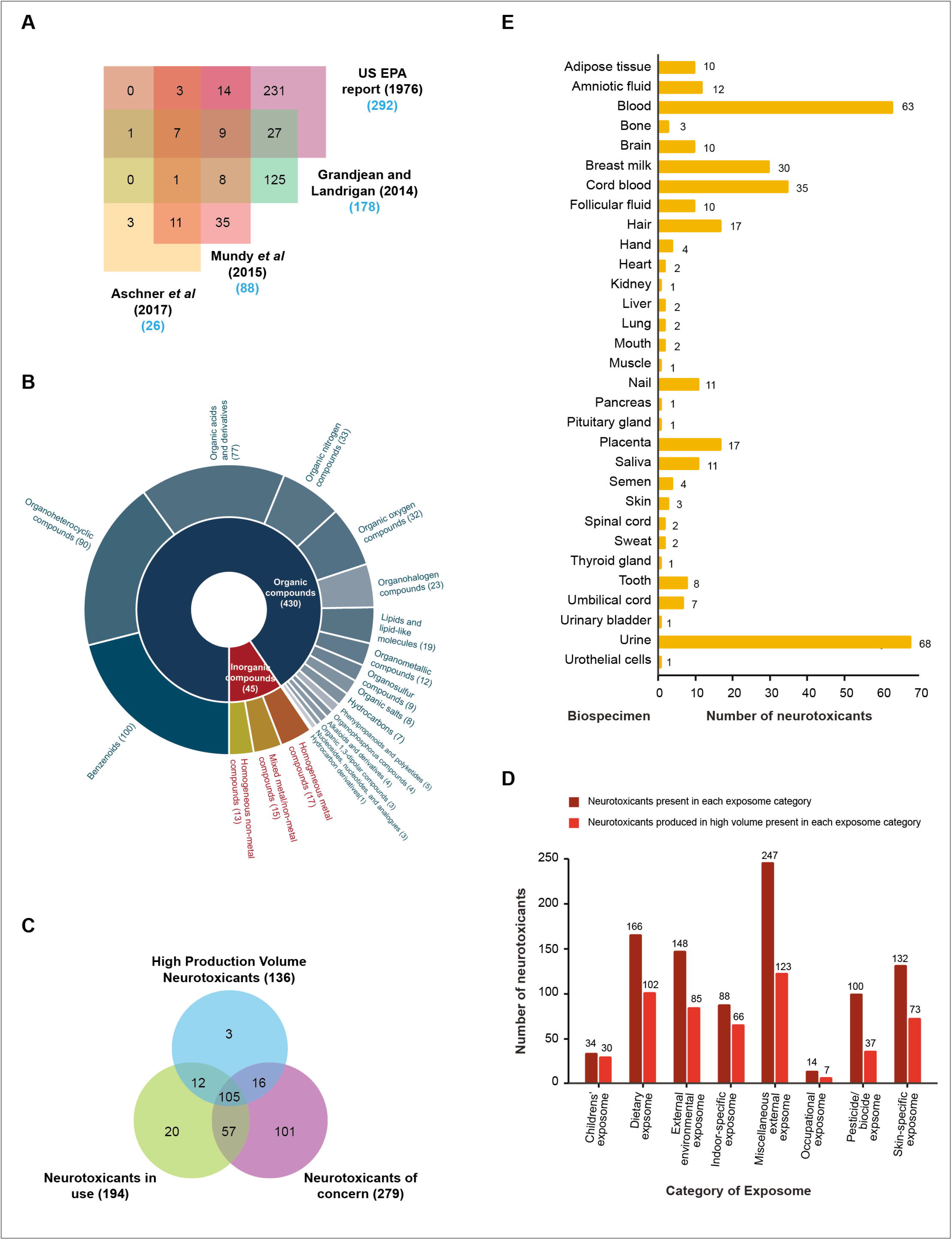
**(A)** Venn diagram showing the occurrence of the 475 potential neurotoxicants compiled in NeurotoxKb across four existing resources, namely, the US EPA report (1976), Grandjean and Landringan (2014), Mundy *et al* (2015), and Aschner *et al* (2017). **(B)** Sunburst plot showing the hierarchal classification of the 475 potential neurotoxicants into 2 chemical kingdoms and 20 chemical superclasses. The number of potential neurotoxicants in each kingdom or superclass is indicated within parenthesis. **(C)** Venn diagram showing the overlap between the sets of potential neurotoxicants present in Substances in use (SIU) lists, Substances of concern (SOC) lists, and High production volume (HPV) lists. Here, the potential neurotoxicants present in SIU lists and SOC lists are labeled as ‘Neurotoxicants in use’ and ‘Neurotoxicants of concern’, respectively. **(D)** Presence of the 475 potential neurotoxicants across chemical lists categorized into 8 exposome categories, namely, Childrens’ exposome, Dietary exposome, External environmental exposome, Indoor-specific exposome, Miscellaneous external exposome, Occupational exposome, Pesticide/biocide exposome, and Skin-specific exposome. This plot displays two bars for each exposome category wherein one bar gives the number of neurotoxicants present in that exposome while other bar gives the number of neurotoxicants that are produced in high volume present in that exposome. **(E)** The bar chart shows the occurrence of the 475 potential neurotoxicants in NeurotoxKb across 31 different human biospecimens.

A unique feature of our knowledgebase NeurotoxKb is the compilation and standardization of the observed neurotoxic effects for the 475 potential neurotoxicants from 835 published studies specific to mammals (Supplementary Table S2). As described in the Methods section, we have undertaken an extensive manual curation effort to compile and standardize the observed neurotoxic effects using MeSH [27] terms for the 475 potential neurotoxicants in NeurotoxKb from associated literature spanning 835 published articles specific to mammals. Notably, the US EPA report [12] contributes a unique set of 414 out of the 835 published articles (~50%) compiled in NeurotoxKb that provide mammalian-specific evidence on potential neurotoxicants. In short, NeurotoxKb compiles the observed neurotoxic effects specific to mammals for the 475 potential neurotoxicants in terms of 148 standardized neurotoxic endpoints from 835 published articles. For the 475 potential neurotoxicants in NeurotoxKb, Supplementary Table S2 gives the compiled evidence specific to mammals, including observed neurotoxic endpoints as MeSH identifiers from associated 835 published studies.

Table 1 presents a comparison of our resource, NeurotoxKb, with the four existing resources, namely, the US EPA report [12], Grandjean and Landrigan [9], Mundy *et al* [13] and Aschner *et al* [14] on potential neurotoxicants. From this table, it is evident that NeurotoxKb will be a valuable resource for future research and monitoring of neurotoxicants due to several additional features in comparison to existing resources.

Based on their environmental sources, we have classified the 475 potential neurotoxicants in NeurotoxKb into 6 broad categories and 41 sub-categories (Methods; Figure 2). It can be seen that majority of the 475 potential neurotoxicants are in the category ‘Agriculture and Farming’ which is followed by ‘Industry’ (Figure 2). Furthermore, we provide a hierarchical classification of the 475 potential neurotoxicants in NeurotoxKb based on their chemical structure (Figure 5B). Of the 475 potential neurotoxicants, 430 are organic while 45 are inorganic (Methods; Figure 5B). Moreover, majority (100) of the 475 potential neurotoxicants belong to chemical superclass ‘Benzenoids’ (Figure 5B). In addition to environmental sources and chemical classification, NeurotoxKb also compiles information on 2D and 3D chemical structure, physicochemical properties, molecular descriptors, predicted ADMET properties and target human genes for the 475 potential neurotoxicants to facilitate future toxicological research.

NeurotoxKb provides the compiled information on the 475 potential neurotoxicants via a user-friendly web interface (Figure 3). Using the web interface, users can access detailed information on any of the potential neurotoxicants via *search* or *browse* options (Figure 3). Using the *simple search* option, users can search for a specific neurotoxicant using its chemical name or standard chemical identifiers (Figure 3B). In addition to simple search, users can also retrieve potential neurotoxicants using filters based on their *physicochemical* properties or *chemical similarity* with respect to a user-specified query compound (Figure 3C-D). Moreover, users can *browse* the potential neurotoxicants in our resource based on their (a) Environmental source, (b) Chemical classification, (c) Presence in chemical regulation or guideline, and (d) Presence in human biospecimen (Figure 3E-H).

### 3.2. Exploration of potential neurotoxicants in external exposomes via chemical regulations and guidelines

Since exposure to neurotoxicants can cause severe, irreversible damage to humans, it is important to understand the extent to which neurotoxicants are monitored via chemical regulations. In this direction, we have explored the presence of 475 potential neurotoxicants in NeurotoxKb across 55 publicly available chemical lists which include inventories, regulations and guidelines (Figure 4; Supplementary Table S3).

As described in the Methods section, the 55 chemical lists have been classified into two broad categories, namely, Substances in use (SIU) and Substances of concern (SOC) (Figure 4; Supplementary Table S3). We find that 311 potential neurotoxicants in NeurotoxKb are present in at least one of the 55 chemical lists (Supplementary Table S4). Figure 5C shows the distribution of these 311 potential neurotoxicants across SIU, SOC and HPV lists (Methods). Notably, 162 potential neurotoxicants are present in both SIU and SOC lists, and further, 105 of these 162 potential neurotoxicants are also produced in high volume (Figure 5C). Among the 311 potential neurotoxicants present in at least one of the 55 chemical lists, Ethylene oxide is present is maximum number (24) of lists which includes both SIU and SOC lists (Supplementary Table S4). Published literature on Ethylene oxide has clearly documented experimental evidence on its neurotoxicity, and humans are mainly exposed to this neurotoxicant via occupational exposure [50,51].

As described in the Methods section, based on the source or route of human exposure, the 55 chemical lists have also been classified into 8 categories of external exposomes (Figure 4; Supplementary Table S3). Figure 5D displays the presence of the 475 potential neurotoxicants across chemical lists categorized into 8 exposome categories. We find that 166 potential neurotoxicants in NeurotoxKb are present in the dietary exposome, specifically as food additives, food packaging materials and food contact substances (Figure 5D). For example, the Pew list of food additives (L7) contains 88 potential neurotoxicants (Figure 4; Supplementary Table S4). Further analysis of the SIU lists classified as Indoor-specific exposome, Pesticide/biocide exposome, Skin-specific exposome or Miscellaneous external exposome found the presence of several potential neurotoxicants compiled in NeurotoxKb (Supplementary Table S4). In other words, we find that several potential neurotoxicants compiled in NeurotoxKb are in regular use. An analysis of the SOC lists classified as Childrens’ exposome, Occupational exposome, Pesticide/biocide exposome, Skin-specific exposome, External environmental exposome or Miscellaneous external exposome found that several potential neurotoxicants compiled in NeurotoxKb are also subject to chemical regulations worldwide.

To highlight the possible implications from this exploratory analysis of the presence of potential neurotoxicants across 55 chemical lists including inventories, regulations and guidelines, we next focus on chemical lists classified into a single category of external exposome, namely, Childrens’ exposome. As neurotoxicants can cause permanent or irreversible damage to neuronal systems [2,7], it is important to monitor and regulate their developmental exposure to children. For this focused analysis, we considered 4 SOC lists namely, Chemicals of concern in plastic toys (L22), Danish EPA Sensitizing Fragrances in Childrens’ Articles (L23), EU Toy Safety Directive (L24), and Washington State Childrens’ Safe Product Act (L25), which contain chemicals prohibited or restricted in children related consumer products. We find that 34 potential neurotoxicants compiled in NeurotoxKb are present in the lists pertaining to Childrens’ exposome, and of these, 30 potential neurotoxicants are also produced in high volume as they are present in HPV lists (Supplementary Table S4). Our observations are indicative of the extent to which these chemicals have been, or are currently being used, in children related products. These 30 potential neurotoxicants warrant further attention, and dedicated monitoring strategies to prevent exposure of children (Supplementary Table S4).

### 3.3. Exploration of potential neurotoxicants in external exposomes via human biospecimens

We next explored the presence of the 475 potential neurotoxicants in NeurotoxKb across human biospecimens compiled from two resources, Exposome-Explorer [16] and CTD [18] (Methods). We find that 91 potential neurotoxicants were detected in at least one of the 31 human biospecimens (Figure 5E; Supplementary Table S5). Among the 91 potential neurotoxicants detected in human biospecimens, Arsenic was detected in maximum number (16) of human biospecimens. Among the 31 human biospecimens, the 68 and 63 potential neurotoxicants were detected in urine and blood, respectively (Figure 5E).

Human fetus is vulnerable to hazardous chemicals such as neurotoxicants [8,9]. Several potential neurotoxicants were detected in human biospecimens related to fetal development or pregnancy. Specifically, we find that 35, 17, 12 and 7 potential neurotoxicants were detected in Cord blood, Placenta, Amniotic fluid and Umbilical cord, respectively (Figure 5E; Supplementary Table S5). Moreover, 30 potential neurotoxicants were also detected in Breast milk via which breastfed infants can be exposed to such chemicals (Figure 5E; Supplementary Table S5). Human brain is sensitive to neurotoxicants and the blood-brain barrier provides only partial protection against such chemicals [52]. We find that 10 potential neurotoxicants were detected in brain (Figure 5E; Supplementary Table S5).

We would like to highlight that well-known neurotoxicants including heavy metals such as Arsenic, Cadmium, Lead, Mercury, Nickel and Selenium, and Perfluoroalkyl substances such as Perfluorooctanesulfonic acid and Perfluorooctanoic acid, were detected in biospecimens related to fetal development, breast milk and brain. These observations underscore the omnipresence of well-known neurotoxicants in our environment, and invite further research and regular monitoring of these chemicals in daily use products and human exposome.

### 3.4. Prioritization of potential environmental neurotoxicants

An exploration of the current chemical regulations and guidelines enabled us to better understand the route and likelihood of human exposure to potential neurotoxicants in their lifetime. We next decided to explore the utility of our resource NeurotoxKb in aiding prioritization of potential neurotoxicants. For this purpose, we have analyzed the presence of the 475 potential neurotoxicants compiled in NeurotoxKb across following lists:

(a) Two lists of high production volume (HPV) chemicals, namely, the USHPV database and the OECD HPV list (Methods). These lists enable to identify potential neurotoxicants that are extensively manufactured, and thus, have a high likelihood of human exposure.
(b) List of substances of very high concern (SVHC) under Registration, Evaluation, Authorisation and Restriction of Chemicals (REACH) regulation of the European Union (EU). SVHC includes chemicals based on their potential to be: (i) Carcinogenic, Mutagenic, toxic to Reproduction (CMR), (ii) disruptive to the endocrine system, (iii) Persistent, Bioaccumulative and Toxic (PBT), and (iv) very Persistent and very Bioaccumulative (vPvB).

Table 2 gives the list of 18 potential neurotoxicants in NeurotoxKb that are also present in both HPV and SVHC lists. Being registered as SVHC, these 18 chemicals are monitored and phased out where necessary, under stringent controls in the EU. These 18 chemicals are associated with multiple types of toxicity (Table 2). Overall, our analysis suggests the need for dedicated monitoring and worldwide prioritization of these 18 potential neurotoxicants. We remark that our analysis of the potential neurotoxicants produced in high volume is limited to HPV lists pertaining to EU and USA due to lack of publicly available HPV lists for other countries. Regulatory bodies in other countries seeking to improve the prioritization of potential neurotoxicants can analyze NeurotoxKb in conjunction with country-specific data on chemical production volume and scale of use.

**Table 2:** List of 18 potential neurotoxicants in NeurotoxKb suggested for prioritization. These 18 chemicals are considered to be substance of very high concern (SVHC) under REACH regulation, and moreover, are present in two lists of high production volume (HPV) chemicals, namely, United States High Production Volume (USHPV) database and Organisation for Economic Co-operation and Development High Production Volume (OECD HPV) list.

| Potential Neurotoxicant | Presence in HPV lists |  | Presence in SVHC list |  |
| --- | --- | --- | --- | --- |
|  | USHPV | OECD HPV | Presence | Criteria |
| Tributyltin oxide | Yes | Yes | Yes | PBT (Article 57d) |
| Lead | Yes | Yes | Yes | Toxic for reproduction (Article 57c) |
| N,N-Dimethylformamide | Yes | Yes | Yes | Toxic for reproduction (Article 57c) |
| Tetraethyllead | Yes | Yes | Yes | Toxic for reproduction (Article 57c) |
| Trichloroethylene | Yes | Yes | Yes | Carcinogenic (Article 57a) |
| Dinoseb | Yes | Yes | Yes | Toxic for reproduction (Article 57c) |
| Nitrobenzene | Yes | Yes | Yes | Toxic for reproduction (Article 57c) |
| Boric acid | Yes | Yes | Yes | Toxic for reproduction (Article 57c) |
| 1-Bromopropane | Yes | Yes | Yes | Toxic for reproduction (Article 57c) |
| 2-Methoxyethanol | Yes | Yes | Yes | Toxic for reproduction (Article 57c) |
| 2,4-Dinitrotoluene | Yes | Yes | Yes | Carcinogenic (Article 57a) |
| Hydrazine | Yes | Yes | Yes | Carcinogenic (Article 57a) |
| Cadmium | Yes | Yes | Yes | Carcinogenic (Article 57a); Specific target organ toxicity after repeated exposure (Article 57(f) - human health) |
| Dibutyl phthalate | Yes | Yes | Yes | Toxic for reproduction (Article 57c); Endocrine disrupting properties (Article 57(f) - human health) |
| Propylene oxide | Yes | Yes | Yes | Carcinogenic (Article 57a); Mutagenic (Article 57b) |
| Acrylamide | Yes | Yes | Yes | Carcinogenic (Article 57a); Mutagenic (Article 57b) |
| Bisphenol A | Yes | Yes | Yes | Toxic for reproduction (Article 57c); Endocrine disrupting properties (Article 57(f) - environment); Endocrine disrupting properties (Article 57(f) - human health) |
| Bis(2-ethylhexyl) phthalate | Yes | Yes | Yes | Toxic for reproduction (Article 57c); Endocrine disrupting properties (Article 57(f) - environment); Endocrine disrupting properties (Article 57(f) - human health) |

A common plasticizer, Bis(2-ethylhexyl) phthalate, is among the 18 potential neurotoxicants suggested for prioritization in this study. Bis(2-ethylhexyl) phthalate, also known as diethylhexyl phthalate or DEHP, is present in 22 out of the 55 chemical lists, of which 7 are SIU lists and 15 are SOC lists (Supplementary Table S4). These 22 chemical lists fall into 6 external exposome categories, namely, Childrens’ exposome, Dietary exposome, External environmental exposome, Indoor-specific exposome, Skin-specific exposome and Miscellaneous external exposome. Bis(2-ethylhexyl) phthalate has been found to impair learning and memory, and cause brain tissue damage in rodents and humans [53,54]. In sum, our resource can aid and offer direction to monitoring organizations and regulatory agencies in identifying, prioritizing and improving the regulations around neurotoxicants.

### 3.5. Interaction of environmental neurotoxicants with neuroreceptors

Neurotoxicants affect the nervous system via diverse molecular mechanisms. Among them two important mechanisms are interference in receptor-mediated processes and modulation of the levels of neurotransmitters by neurotoxicants [6]. We have used ToxCast [36] database to obtain the target human genes of the 475 potential neurotoxicants in NeurotoxKb (Methods). Based on human-specific assays in ToxCast [36], we were able to obtain 255 target human genes for 220 out of the 475 potential neurotoxicants in NeurotoxKb (Supplementary Table S6). Further investigation of the 255 target human genes of the 220 potential neurotoxicants revealed that 27 target genes correspond to neuroreceptors. We find that 38 potential neurotoxicants in NeurotoxKb target at least one of these 27 neuroreceptors (Figure 6; Supplementary Table S6). Among these 38 potential neurotoxicants, 4 neurotoxicants namely, Mercuric chloride, Haloperidol, Triphenyltin hydroxide and Perfluorooctanesulfonic acid (PFOS), target 10 or more neuroreceptors (Figure 6). Among the 27 neuroreceptors which are targets of at least one potential neurotoxicant, the neuroreceptor OPRM1 (Opioid Receptor Mu 1) for endogenous opioids such as β-endorphin and endomorphin, was found to interact with 15 potential neurotoxicants. Other neuroreceptors which are targets of at least 10 potential neurotoxicants include the receptor DRD1 (Dopamine receptor D1) for neurotransmitter dopamine, and the receptors HTR6 (5-Hydroxytryptamine Receptor 6) and HTR7 (5-Hydroxytryptamine Receptor 7) for the neurotransmitter serotonin (Figure 6; Supplementary Table S6). In future, an in depth analysis of chemical-gene interactions will shed new insights on the molecular mechanisms via which the exposure to the 475 potential neurotoxicants in NeurotoxKb can lead to documented neurotoxic endpoints in mammals.

**Figure 6:**
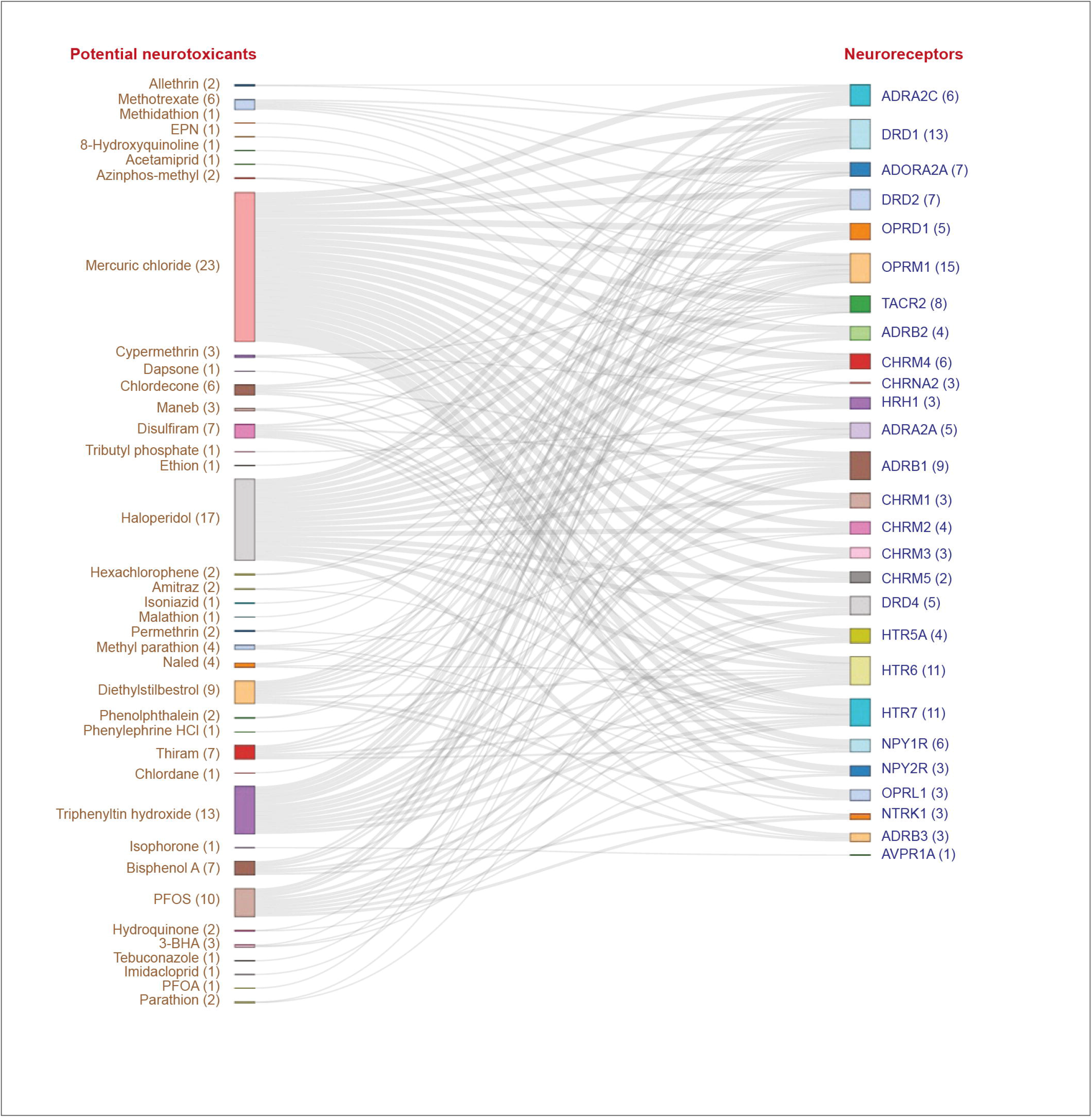
The bipartite network of 38 potential neurotoxicants in NeurotoxKb that target 27 human neuroreceptors. Besides each potential neurotoxicant, the number of target neuroreceptors is indicated within parenthesis. Besides each neuroreceptor, the number of potential neurotoxicants targeting it is indicated within parenthesis.

### 3.6. Network based visualization of the chemical space of environmental neurotoxicants

Recently, chemical similarity approaches have received due attention in early prediction of chemical risk [21,22]. In this direction, we have constructed a chemical similarity network (CSN) for the 475 potential neurotoxicants compiled in NeurotoxKb (Methods; Figure 7). In Figure 7, it is seen that the CSN of 475 potential neurotoxicants is fragmented into 60 connected components with number of neurotoxicants ≥ 2 and 286 isolated neurotoxicants. Moreover, the largest connected component consists of only 13 potential neurotoxicants (Figure 7). In Figure 7, we have coloured the nodes based on the number of aromatic rings in the corresponding neurotoxicant. It can be seen that neurotoxicants belonging to a connected component typically have the same number of aromatic rings. Altogether, this preliminary analysis of the CSN of potential neurotoxicants reveals a fragmented network, and thus, the associated toxicological space has high chemical diversity.

**Figure 7:**
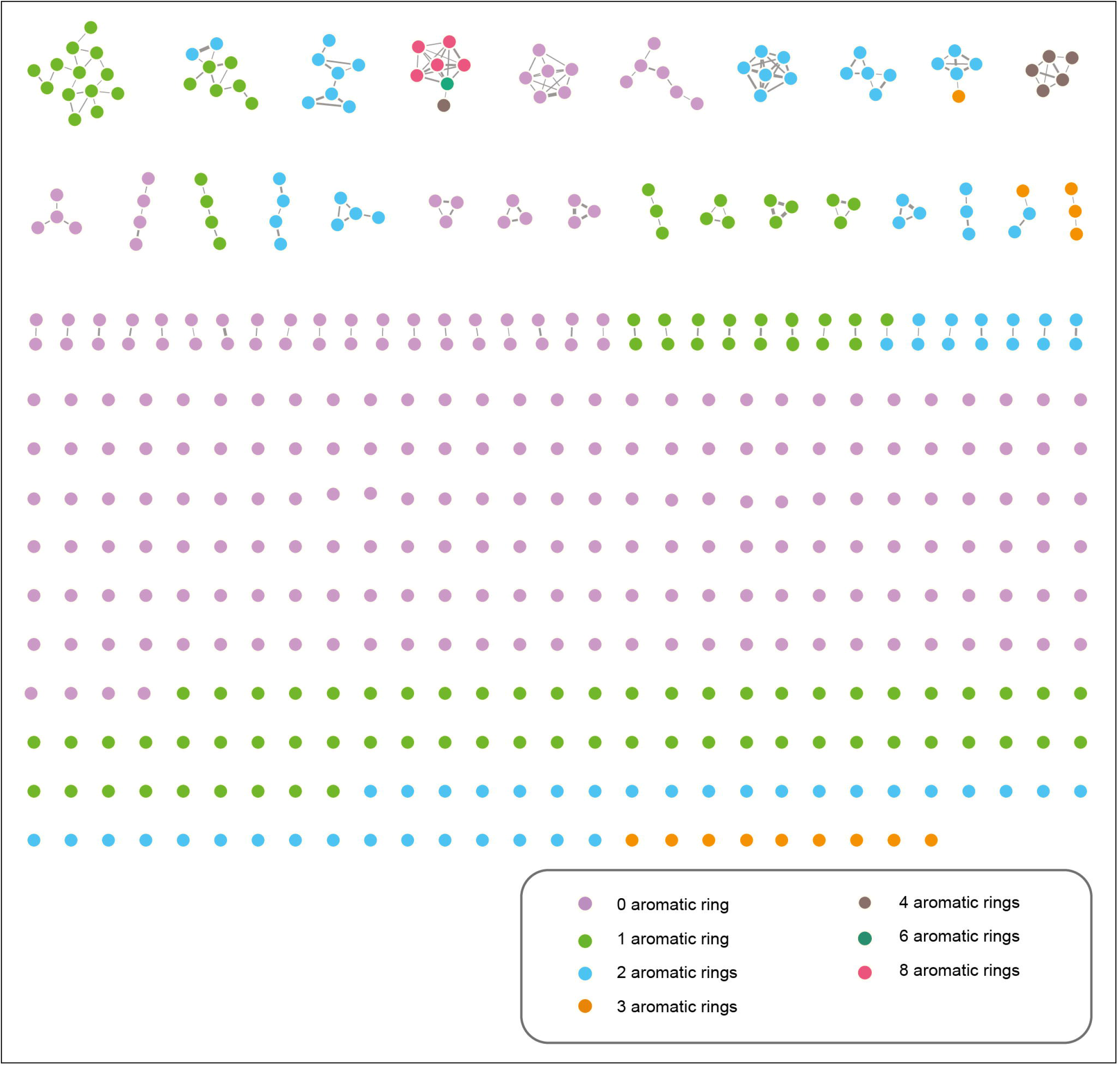
Chemical similarity network (CSN) of the 475 potential neurotoxicants in NeurotoxKb. In this figure, there are 475 nodes corresponding to the 475 potential neurotoxicants, and there is an edge between any pair of nodes if the corresponding Tanimoto coefficient (Tc) value is ≥ 0.5. Further, nodes are coloured based on the number of aromatic rings present in the corresponding neurotoxicants, while the thickness of the edges indicate the Tc value between the corresponding neurotoxicants. Here, the connected components of the CSN are displayed in the decreasing order of the number of nodes in each component.

## 4. Conclusions

The Swiss philosopher and poet, Henri-Frédéric Amiel (1821-1881), once stated that: “To repair is twenty times more difficult than to prevent”. The quote is apt for the management of hazardous chemicals including environmental neurotoxicants. Since neurotoxicants can cause permanent or irreversible damage to the nervous system [7,2,3], early screening of environmental chemicals with potential to cause neurotoxicity is important for human well-being. In this direction, a comprehensive resource on potential neurotoxicants compiling published evidence specific to mammals, can aid in monitoring and regulation of human neurotoxicants. Here, we present such a comprehensive resource, NeurotoxKb, with compiled information on 475 potential non-biogenic neurotoxicants curated from 835 published studies specific to mammals. The entire compiled information on the 475 potential neurotoxicants in NeurotoxKb can be easily accessed and retrieved via a user-friendly and interactive web interface.

Humans are exposed to environmental neurotoxicants via diverse sources (Figure 2). Firstly, a comparative analysis of NeurotoxKb and 55 chemical lists which include inventories, regulations and guidelines, found that several potential neurotoxicants are both in regular use and produced in high volume (Figures 4 and 5C). Secondly, a comparative analysis of NeurotoxKb and chemicals detected in 31 different human biospecimens, found that several potential neurotoxicants have been detected in different biospecimens (Figure 5E). In other words, our comparative analysis with chemicals in regulatory lists or those detected in human biospecimens confirm the omnipresence of potential neurotoxicants in different categories of external exposomes (Figure 4). Furthermore, based on a comparative analysis of NeurotoxKb with SVHC REACH regulation and HPV chemicals, we present a hazard priority list of 18 potential neurotoxicants (Table 2). In sum, NeurotoxKb can be used for identification and prioritization of environmental neurotoxicants in human exposomes.

A unique feature of our resource on potential neurotoxicants is the compilation and standardization of neurotoxic endpoints from published studies specific to mammals. In future, it will be worthwhile to leverage this compiled information in NeurotoxKb to develop adverse outcome pathways [55] for different neurotoxicants. We envisage that such an extension of our knowledgebase can further aid risk assessment of environmental chemicals.

### CRediT author contribution statement

**Janani Ravichandran:** Conceptualization, Data curation, Formal analysis, Software, Visualization, Writing – original draft. **Bagavathy Shanmugam Karthikeyan:** Conceptualization, Data curation, Formal analysis, Writing – original draft. **Palak Singla:** Formal analysis, Data curation. **S. R. Aparna:** Writing – original draft. **Areejit Samal:** Conceptualization, Investigation, Formal analysis, Supervision, Writing – review & editing.

## Acknowledgements

We thank B. Raveendra Reddy for help in setting up the webserver and Murugesan Karuppasamy for assistance in data compilation. AS acknowledges support from the Science and Engineering Research Board (SERB) India through a Ramanujan fellowship (SB/S2/RJN-006/2014), the Department of Atomic Energy (DAE) India, and the Max Planck Society Germany through a Max Planck Partner Group in Mathematical Biology. The funders have no role in study design, data collection, data analysis, manuscript preparation or decision to publish.

## Supplementary Tables

**Table S1:** List of 475 potential neurotoxicants in NeurotoxKb along with their CAS and Pubchem identifiers compiled from four existing resources, namely, the US EPA report (1976), Grandjean and Landrigan (2014), Mundy *et al* (2015), and Aschner *et al* (2017). Of the 475 potential neurotoxicants, 292, 178, 88 and 26 were compiled from the US EPA report (1976), Grandjean and Landrigan (2014), Mundy *et al* (2015), and Aschner *et al* (2017), respectively.

**Table S2:** List of observed neurotoxic endpoints specific to mammals for the 475 potential neurotoxicants in NeurotoxKb compiled from published literature. For the potential neurotoxicants listed in the US EPA report (1976), Mundy *et al* (2015), and Aschner *et al* (2017), the neurotoxic endpoints were compiled from the associated literature reported in the respective list. The neurotoxic endpoints for the potential neurotoxicants listed in Grandjean and Landrigan (2014) were compiled from the Hazardous Substances Data Bank (HSDB). The compiled neurotoxic endpoints were unified into standardized MeSH terms and identifiers.

**Table S3:** The table provides a description of the publicly available 55 chemical lists that are a part of chemical inventories, regulations and guidelines. These lists were broadly classified into ‘Substances in use (SIU)’ and ‘Substances of concern (SOC)’ based on the nature of substances. Further, they have been categorized into Childrens’ exposome, Dietary exposome, External environmental exposome, Indoor-specific exposome, Miscellaneous external exposome, Occupational exposome, Pesticide/biocide exposome, and Skin-specific exposome based on the route or source of exposure.

**Table S4:** The table lists the 311 potential neurotoxicants in NeurotoxKb that are present in at least one of the 55 chemical lists that are part of inventories, regulations and guidelines. Of these 311 potential neurotoxicants, 133 were found to be produced in high volume.

**Table S5:** List of 91 potential neurotoxicants in NeurotoxKb detected across 31 different human biospecimens compiled from Exposome-Explorer and CTD.

**Table S6:** List of target human genes for the 220 potential neurotoxicants in NeurotoxKb, based on 545 ToxCast assay component endpoints specific to human. Of the 220 potential neurotoxicants, 38 were found to target at least one of 27 neuroreceptors that are known to mediate neuronal responses.

